# An epithelial-mesenchymal-amoeboid transition gene signature reveals subtypes of breast cancer progression and metastasis

**DOI:** 10.1101/219410

**Authors:** Amin Emad, Tania Ray, Tor W. Jensen, Meera Parat, Rachael Natrajan, Saurabh Sinha, Partha S. Ray

**Author notes:** Joint senior authors. Corresponding Authors: Partha S. Ray Onconostic Technologies, Inc. 60 Hazelwood Drive, Suite 208 Champaign, IL, USA 61820 Saurabh Sinha 2122 Siebel Center 201 N. Goodwin Ave Urbana, IL, USA 61801.

## Abstract

Cancer cells within a tumor are known to display varying degrees of metastatic propensity but the molecular basis underlying such heterogeneity remains unclear. We analyzed genome-wide gene expression data obtained from primary tumors of lymph node-negative breast cancer patients using a novel metastasis biology-based Epithelial-Mesenchymal-Amoeboid Transition (EMAT) gene signature, and identified subtypes associated with distinct prognostic profiles. EMAT subtype status improved prognosis accuracy of clinical parameters and statistically outperformed traditional breast cancer intrinsic subtypes even after adjusting for treatment variables. Additionally, analysis of 3D spheroids from an in vitro isogenic model of breast cancer progression reveals that EMAT subtypes display progression from premalignant to malignant and pre-invasive to invasive cancer. EMAT classification is a biologically informed method to assess metastasis risk in early stage, lymph node-negative breast cancer patients.

## INTRODUCTION

Metastasis accounts for nearly 90% of cancer related mortality, and cancer cells within a tumor are known to possess different metastatic potentials (Fidler & Kripke, 1977). However, the molecular basis underlying the observed heterogeneity in metastatic proclivity remains unclear and a suitable molecular classification is lacking. Intrinsic molecular subtypes of breast cancer have been associated with distinct metastatic predilections for one organ or the other, but are not necessarily associated with an increased metastatic propensity per se (Smid, Wang et al., 2008). For instance, an intrinsic subtype that displays a higher rate of brain metastasis does not necessarily mean all patients, or even the majority of patients diagnosed with that subtype of cancer will go on to manifest with metastatic disease in the brain. Clearly other factors, independent of and in addition to those that determine the intrinsic molecular subtype of the cancer, influence its invasive potential and metastatic propensity.

Although implicated in cancer progression and metastasis, the clinical significance of processes like epithelial-to-mesenchymal transition (EMT) and mesenchymal-to-amoeboid transition (MAT) remains to be fully appreciated. EMT, a cellular transformation process that plays a key role in embryonic development, is widely considered to be one such factor influencing metastasis. Cancer cells derepress the normally silenced EMT molecular program, acquiring malignant properties that enable them to invade tissues surrounding their site of origin thereby effectively spreading and colonizing distant sites (Thompson, Paik et al., 1992, Yang & Weinberg, 2008). Likewise, MAT is another process that plays an important role in embryonic development and is similarly reawakened by cancers during the invasion-metastasis cascade (Wolf & Friedl, 2006, Wolf, Mazo et al., 2003).

Since the EMT process is exploited by cancer cells progressing to metastasis, there have been several attempts to subtype patient tumors based on an EMT signature, but these have not been successful in demonstrating a discernible difference in associated breast cancer prognosis (Marsan, Van den Eynden et al., 2014, Tan, Miow et al., 2014, Taube, Herschkowitz et al., 2010). Additionally, for cancer cells that have already transitioned through EMT but are facing microenvironmental (e.g. hypoxia) or xenobiotic (e.g. chemotherapy) stress, MAT may be an effective adaptive response to bypass the stress (Lehmann, te Boekhorst et al., 2017). Indeed a recent report of effectively thwarting metastatic spread through simultaneous targeting of both mesenchymal and amoeboid motility in an animal model of cancer progression appears to support this notion (Jones, Kelley et al., 2017). We thus hypothesized that the true clinical and prognostic significance of EMT as a driving process in cancer progression towards distant metastasis cannot be fully appreciated unless it is considered in the context of being complemented by the conditional occurrence of MAT as well. Only when both processes are considered to coexist and undergo plastic interchange triggered by environmental pressures can the clinical significance of both be recognized and prognostic impact demonstrated. We therefore sought to develop a more inclusive gene expression signature that accurately captures EMT, MAT, and the variable dynamic co-occurrence of both the processes in the same tumor. In this study our aims were to i) elucidate prognostic subtypes in a primary tumor based on an EMT-MAT continuum that captures the heterogeneity of metastatic propensity and ii) to more comprehensively define biologically informed subtypes predictive of breast cancer metastasis and survival.

We constructed a gene signature (henceforth referred to as the EMAT signature) by combining a previously reported signature of EMT (Taube et al., 2010), derived from gene expression data of multiple distinct EMT-inducing perturbation experiments using cancer cells, with a newly generated MAT signature, derived following identical methodology to minimize derivation bias. This gene signature was derived purely from cancer cells as our objective was to solely gauge the potential contribution of cancer cells in mediating metastasis independent of the contribution of other cells in the tumor microenvironment or that of the tumor stroma. We utilized this new EMAT gene signature to probe lymph node-negative human primary breast cancer gene expression datasets and were able to delineate informative subtypes of breast cancer that possess distinct molecular features, morphology/motility phenotypes and metastatic propensities. While EMT or MAT signatures on their own could not identify clusters of distinct prognosis, their combined consideration as EMAT subtypes significantly improved survival prediction of standard clinical parameters and significantly outperformed prognosis accuracy of other known subtypes of breast cancer even when adjusting for treatment variables. Focusing on lymph node-negative samples allowed us to identify clusters that can predict clinical outcome at an early clinical stage of cancer. In addition, since cancer cells may utilize different mechanisms to promote metastasis before and after lymph node invasion, we studied lymph node-negative and positive samples separately and found the clinical characteristics of EMAT-defined subtypes to be largely similar, with some key differences. Furthermore, we identified transcription factors that may play key regulatory roles in establishing the newly identified EMAT subtypes and may be used as potential biomarkers for these subtypes. Finally, the identified subtypes were validated in a comparable but independent, lymph node negative, treatment-naïve dataset and their enrichment was examined in an established and accepted *in vitro* cell line 3D spheroid model of molecular drivers of breast cancer progression, further confirming that EMAT subtypes reflect the progressive cellular transition from benign to malignant and from non-invasive to invasive cancer. Our successful demonstration of prognostic stratification of breast cancer patients based on an EMAT signature opens up exciting avenues for future research into the mechanisms of the invasion/metastasis cascade.

## RESULTS

### An EMAT gene signature predicts clinical outcome in breast cancer patients

We constructed an “EMAT” gene signature comprising a list of 253 previously reported EMT-related genes (Taube et al., 2010) and 138 newly derived MAT-related genes obtained through analysis of publicly available gene expression data from multiple distinct MAT-inducing cell perturbation experiments (Taddei, Giannoni et al., 2014) (see Methods and Supplementary Table S1). The analyzed dataset comprised mRNA gene expression profiles and clinical data corresponding to 562 lymph node-negative (LN) breast cancer primary tumors from the METABRIC study (Curtis, Shah et al., 2012). (We used lymph node negativity as a criterion to ensure that the samples were obtained during early clinical stages of the invasion-metastasis cascade.) We performed hierarchical clustering of samples (see Methods), represented by expression levels of EMAT genes (Figure 1A), evaluated resulting partitions with n = 3, 4, or 5 clusters, and found that n = 4 yields the best grouping of samples based on average silhouette scores (data not shown) (Rousseeuw, 1987). This clustering (Supplementary Table S2) provided a clear separation of the Kaplan-Meier survival curves (p = 2.42E-4, log rank test, Figure 1C), even though the clustering procedure was not privy to survival data. In addition, univariable and multivariable Cox regression analysis (when considering clinical parameters and treatment variables) showed that these clusters provide a statistically significant prognostic value (Supplementary Table S3). We refer to the four clusters, in decreasing order of their 10-year disease-specific survival (DSS) probabilities, as EMAT1, EMAT2, EMAT3 and EMAT4. Interestingly, clusters EMAT2 and EMAT3 have very similar Kaplan-Meier survival curves (Figure 1C), even though their gene expression profiles are distinct (indicated by orange bar in Figure 1A).

**Figure 1:**
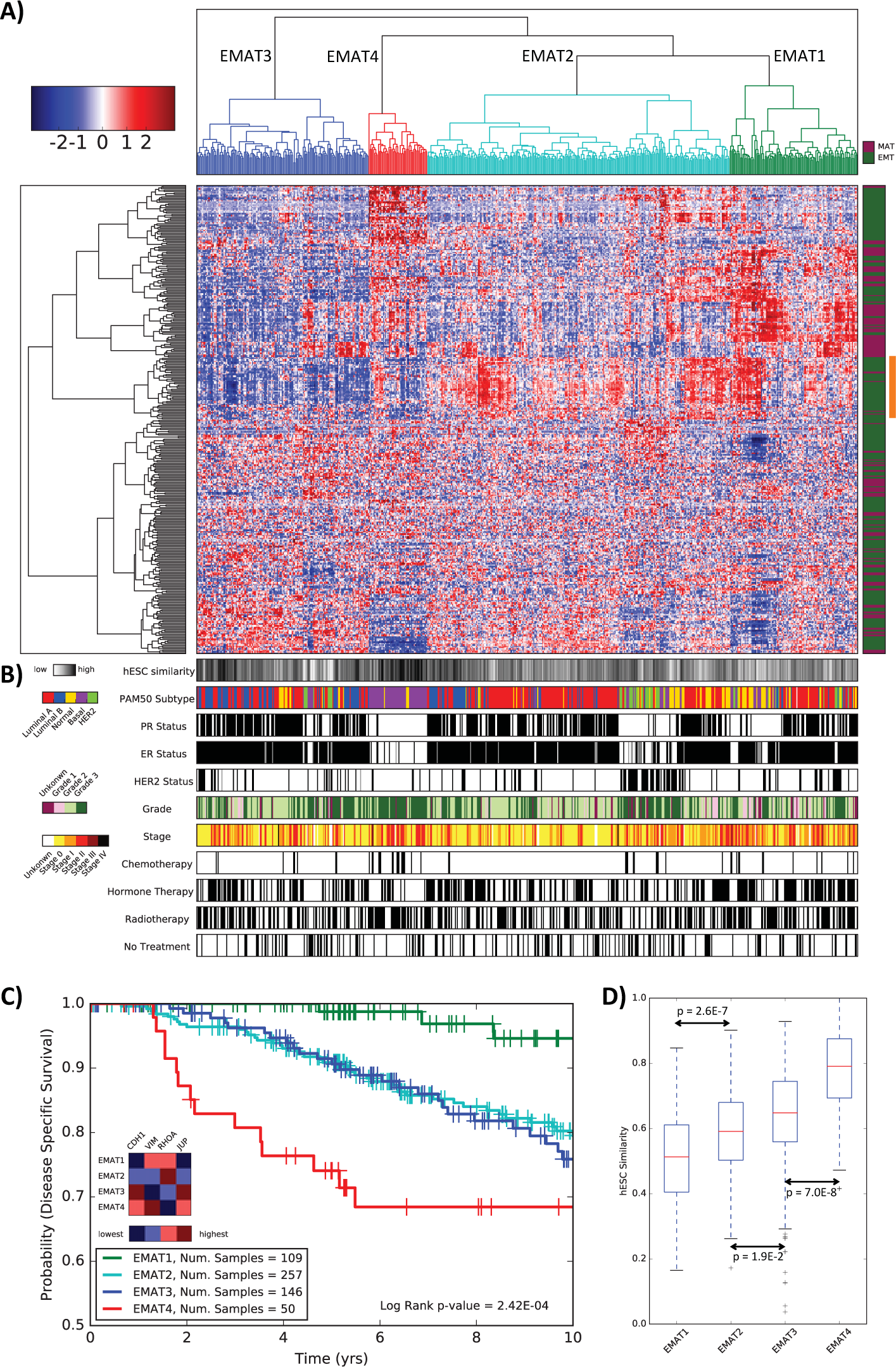
EMAT clusters and their characteristics. (A) EMAT clusters based on lymph node-negative METABRIC samples obtained using hierarchical clustering. The heatmap shows the normalized expression of EMAT genes (rows) in each sample (columns). Sample dendogram colors are chosen to match those of Kaplan-Meier plots in Figure 1C. (B) Characterization of samples based on similarity to hESC, PAM50 subtypes, ER, PR and HER2 status, stage, grade, and type of treatment. Spearman’s rank correlation, scaled between 0 and 1 using min-max normalization, is used as the measure of similarity of samples to hESC, in which 0 and 1 represent least similar and most similar, respectively. (C) Kaplan-Meier plots corresponding to n = 4 clusters. The heatmap shows the relative ranking of the average expression of four biomarkers in each cluster compared to other clusters. (D) The box plots show the distribution of hESC similarity of the samples in each cluster. The similarity is defined as the Spearman’s rank correlation (scaled between 0 and 1) between expression profiles of H1 hESC lines and each sample. The p-values (calculated using a one-sided t-test) show how significant the differences between two adjacent EMAT clusters are with respect to their similarity to hESC. The significance p-value for the cluster with the least similarity to hESC (EMAT1) and the cluster with the most similarity to hESC (EMAT4) is p = 1.7E-23.

To characterize each cluster obtained above, we examined the expression of CDH1, VIM, RHOA, and JUP, well-accepted biomarkers of Epithelial (E), Mesenchymal (M), Amoeboid (A), and collective cell migration morphologies in breast cancer (Weber, Bjerke et al., 2012), respectively. We assessed the expression of JUP as a collective cell migration marker because while EMT and MAT usually manifest as single cell migration, both epithelial and amoeboid collective cell migration have also been observed to contribute to the metastatic spread of cancer cells (Friedl & Wolf, 2008). EMAT4, the cluster with the lowest DSS probability, showed M-like characteristics (VIM was over-expressed in this cluster compared to the other three clusters, p = 4.8E-8, unpaired two-tailed t-test). Cluster EMAT3, had the lowest average expression of VIM (p = 3.7E-79) and the highest average expression of CDH1 (p = 1.3 E-3) and JUP (p = 8.3E-6), suggesting that early onset of epithelial collective cell migration may be manifested in this group of patients. EMAT2 had the highest average expression of RHOA (p = 3.8E-3), suggesting A-like characteristics. Finally, EMAT1 had high expressions of VIM and RHOA but low expression of CDH1 and JUP, with the average expression of JUP being smallest (p = 3.9E-15) in this cluster. These results indicate existence of clusters having hybrid characteristics rather than discrete E-, M- and A-subtypes and emphasize the advantage of using the EMAT signature over using only E, M, or A biomarkers to distinguish groups of patient tumors associated with distinct prognosis. It is worth mentioning that genes most differentially expressed in each cluster (Bonferroni adjusted p < 0.01, t-test) included both EMT and MAT genes for all clusters (Supplementary Table S4), suggesting the involvement of both signatures in identification of these clusters.

Our finding that an EMAT signature allows grouping of patients with significantly different DSS is particularly interesting in light of previous reports where EMT gene signatures alone failed to do so (Tan et al., 2014, Taube et al., 2010). We confirmed these previous observations in our dataset by clustering the tumor samples based on the expression of the EMT signature (subset of the EMAT signature above) using hierarchical clustering into two clusters, following the method of previous studies. We noted that biomarkers of Epithelial (CDH1) and Mesenchymal (VIM) morphology were differentially expressed between these two clusters, with VIM being over-expressed in EMT1 (p = 1.8E-69) and CDH1 over-expressed in EMT2 (p = 1.4E-2). In spite of this distinction of biomarker expression, Kaplan-Meier analysis (Supplementary Figure S1A) did not show a significant difference between the clinical outcomes of these two groups (p = 0.28). We also clustered samples based on the expression of MAT genes, and noted differential expression of known biomarkers of Mesenchymal (VIM, p = 2.7E-7) and Amoeboid (RHOA, p = 6.4E-4) morphology. However, Kaplan-Meier analysis (Supplementary Figure S1B) did not show a significant difference between the clinical outcomes of these two groups (p = 0.56). It would thus appear that while both EMT and MAT processes have been demonstrated to play contributory roles in the biology of metastasis, when considered individually they cannot completely capture the diversity that exists in the metastatic propensity of breast tumors, as evidenced by their lack of individual prognostic value.

### EMAT clusters provide prognostic information not present in clinical parameters or intrinsic subtypes

Next, we examined whether the EMAT clusters are associated with clinical parameters or previously described breast cancer subtypes (Figure 1B). Of note is the visually apparent enrichment of EMAT4, the cluster with worst survival, with triple negative (negative for ER, PR, HER2) and basal-like (PAM50 subtype (Parker, Mullins et al., 2009), purple) patients. In addition, most HER2 positive patients (green) appear in EMAT2. Similarity to human embryonic stem cells (hESC), representing unicellular and stem-like characteristics, has been previously shown to be associated with elevated metastasis risk (Ben-Porath, Thomson et al., 2008). Figure 1D shows that EMAT clusters display worsening prognosis proportionate to their degree of similarity to hESC, consistent with this theory. This was despite the fact that derivation of the EMAT gene signature was not designed to intentionally enrich for stem cell traits. This provides further evidence in support of the EMAT clusters potentially representing a progressive transition from less stem-like to more stem-like cell states, and less invasive to more invasive modes of cancer.

To assess the quantitative significance of the above associations, we computed the enrichment p-value (hypergeometric test) of each EMAT cluster with respect to tumor size, PAM50 subtypes, and receptor status (Figure 2A-C). While the presence of small tumors in clusters with good prognosis is expected, the enrichment of EMAT1 (the cluster with the best prognosis) in large tumors is particularly interesting, as it suggests that large tumors do not necessarily result in poor survival in the absence of necessary metastatic mechanisms (Comen, Norton et al., 2011). Figure 2B shows that EMAT1 was moderately enriched in the ‘normal-like’ PAM50 subtype (p = 5.59E-3), EMAT2 in Luminal A (p = 1.84e-06), EMAT3 in Luminal B (7.34E-12), and EMAT4 in basal-like subtype (p = 7.73E-41). It is critical to note that in spite of these enrichments, the EMAT clusters are distinct from these molecular subtypes. Figure 2D shows that 68% of normal-like samples are in clusters other than EMAT1, 43% of Luminal A patients are in clusters other than EMAT2, 47% of Luminal B patients are in clusters other than EMAT3 and 46% of Basal patients are in clusters other than EMAT4. In addition, HER2 patients are distributed in EMAT1, EMAT2 and EMAT3 clusters. Finally, EMAT3 was enriched in ER positive (p = 1.76E-06) and PR positive (p = 2E-06) samples, while EMAT4 was enriched in ER negative (p = 2.64E-28), PR negative (p = 4.18E-18), and HER2 negative (p = 4.8E-3) samples (see Table 1 for a summary).

**Figure 2:**
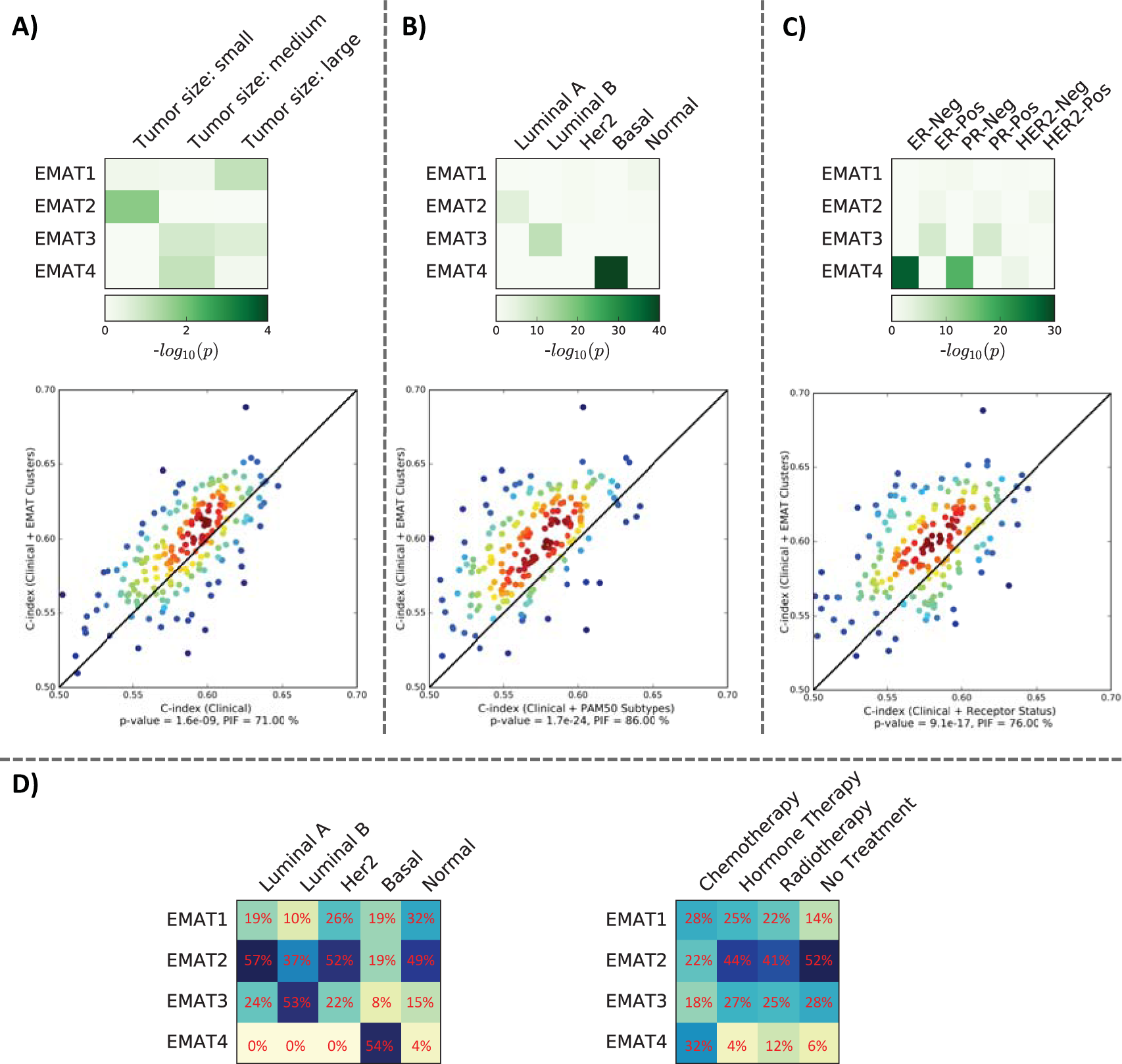
Enrichment of EMAT clusters in other breast cancer subtypes and systematic comparison of their prognostic power using cross-validation. The heatmaps show the -log10 (p-value) of enrichment of EMAT clusters in different subtypes or clinical parameters (using a hypergeometric test). Tumor size less than 2cm is considered small, between 2cm and 5cm is considered medium and larger than 5cm is considered large. The scatter plots compare the performance (measured in C-index) of Cox regression predictions using EMAT cluster status and clinical parameters (y-axis) versus other predictors (x-axis). Here, half of the samples were randomly selected as the training set, and grouped into n = 4 clusters using the EMAT signature. A Cox regression model was trained on these samples, using clinical variables as well as EMAT cluster status. In parallel, Cox regression models were trained on the same samples using the three other types of predictors. Each trained model was then used to estimate the expected survival of the remaining samples (i.e., test samples). In this step, EMAT cluster status was assigned to test samples using a centroid-based classifier trained on the training samples. The p-values were calculated using a one-sided Wilcoxon signed rank test and represent the significance of the improvement obtained using EMAT cluster status and clinical parameters as compared to other predictors. The measure PIF shows the percent of times in which EMAT + clinical parameters provided a more accurate prediction compared to the baseline. (A) The heatmap shows enrichment of EMAT clusters by samples of different tumor sizes. The scatter plot shows performance of Cox regression predictions using EMAT + clinical parameters versus clinical parameters alone. (B) The heatmap shows enrichment of EMAT clusters by samples of different PAM50 molecular subtypes. The scatter plot shows performance of Cox regression predictions using EMAT + clinical parameters versus PAM50 subtypes + clinical parameters. (C) The heatmap shows enrichment of EMAT clusters by samples of different receptor status. The scatter plot shows performance of Cox regression predictions using EMAT + clinical parameters versus receptor status + clinical parameters. (D) The heatmaps show the distribution of patients in each PAM50 subtypes as well as different treatments in EMAT clusters.

**Table 1:**
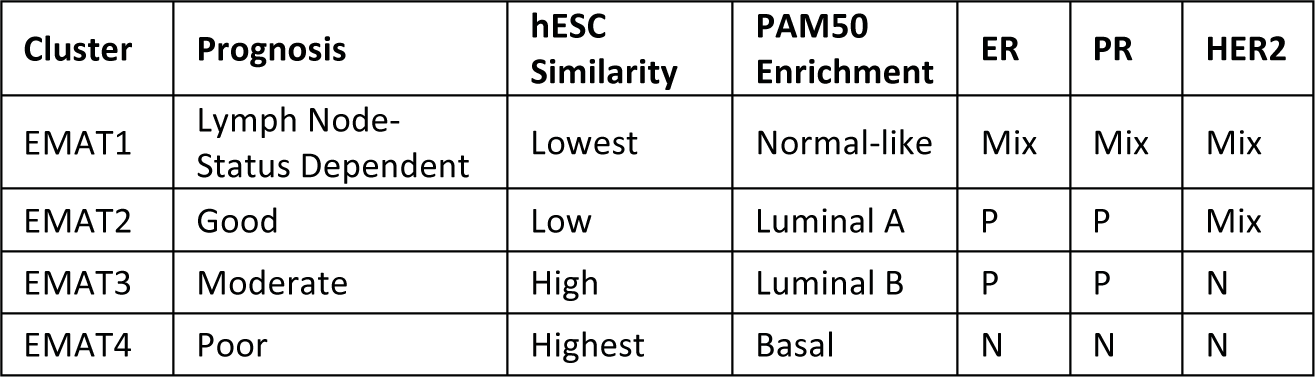
A summary of the characteristics of the EMAT clusters obtained using both lymph node-negative and lymph node-positive breast cancer patients from the METABRIC study. In this table, P stands for positive and N for negative. EMAT1 is a cluster with good prognosis in LN samples but with poor prognosis in LP samples. However, in both cases, it has the least similarity to hESC and is enriched in normal-like PAM50 subtype of breast cancer. EMAT2, the cluster with a relatively good prognosis in both LN and LP samples, has little similarity to hESC, is enriched in Luminal A subtype and in ER-positive and PR-positive samples. EMAT3, the cluster with a relatively moderate prognosis, has a high degree of similarity to hESC, is enriched in Luminal B subtype and in ER-positive, PR-positive and HER2-negative samples. EMAT4, the cluster with the worst prognosis, shows the highest degree of similarity to hESC, is enriched in the basal-like subtype of breast cancer as well as ER-negative, PR-negative and HER2-negative samples.

| Cluster | Prognosis | hESC Similarity | PAM50 Enrichment | ER | PR | HER2 |
| --- | --- | --- | --- | --- | --- | --- |
| EMAT1 | Lymph Node-Status Dependent | Lowest | Normal-like | Mix | Mix | Mix |
| EMAT2 | Good | Low | Luminal A | P | P | Mix |
| EMAT3 | Moderate | High | Luminal B | P | P | N |
| EMAT4 | Poor | Highest | Basal | N | N | N |

Given the observation that EMAT clusters are enriched in intrinsic subtypes defined based on PAM50 and receptor status, we sought to determine whether EMAT clustering could provide prognostic information beyond what is captured by the aforementioned subtypes. To this end, we trained a Cox regression model on the LN samples from the METABRIC dataset using four types of predictors: 1) clinical parameters, including age, tumor size, use of chemotherapy, use of hormone therapy, use of radiotherapy and no treatment, 2) clinical parameters and receptor status, 3) clinical parameters and PAM50 subtype status, and 4) clinical parameters and EMAT cluster status. The Cox regression model trained on clinical and EMAT cluster status had the smallest p-value using a likelihood ratio test (p = 6.6E-5 compared to p = 1.0E-2 for clinical parameters and receptor status, p = 1.5E-2 for clinical parameters and PAM50 subtypes, and p = 4.2E-3 for clinical parameters only). This was a promising result, but not an entirely convincing comparison since EMAT cluster status was defined based on samples used in the Cox regression test, while the other subtypes were defined *a priori*.

To rigorously compare the predictive ability of the above classes of predictors while removing the effect of the varying number of predicting features, and also to test the generalizability of these models on unseen data, we next used a cross-validation framework. In this framework, the samples were randomly divided in two groups of (almost) equal size and a Cox regression model trained on one half was used to estimate the expected survival on the other half. This process was repeated 200 times, each time using a distinct random partition of data. The Cox regression model using EMAT labels along with clinical parameters provided the best predictions (Figure 2A-C bottom panels), evaluated using a one-sided Wilcoxon signed rank test on paired C-index values of the compared methods as well as another measure called percentage of improved folds (PIF) (Emad, Cairns et al., 2017) defined as percent of the partitions in which one class of features outperforms another class.

These results indicate three major points. First, although some of the EMAT clusters are enriched in previously known molecular subtypes of breast cancer (e.g. PAM50), they are quite distinct from these subtypes (Figure 2D). In addition, the EMAT clusters are better predictors of patient survival outcome than PAM50 or receptor-based subtypes. Finally, even though clinical parameters including the type of treatment are important in predicting survival outcome, the EMAT clusters do not simply recover types of treatments given to patients (Figure 2D) but rather provide extra information that are useful in predicting patient prognosis, as is evident from Figure 2A.

### Cross-dataset analysis shows similar survival behavior of the EMAT clusters

We next evaluated the prognostic power of the EMAT clusters on an independent, lymph node negative, treatment-naive dataset. To this end, we developed a subtyping procedure using a centroid-based classifier, trained on the gene expression profiles and EMAT cluster status of the METABRIC LN samples, to assign EMAT “subtypes” to any new dataset (see Methods). We obtained gene expression and clinical data corresponding to lymph node-negative, treatment-naive breast cancer primary tumors from (Schmidt, Böhm et al., 2008) (GEO accession number: GSE11121) and determined their EMAT subtypes. One advantage of this dataset is that it contains distant metastasis-free survival (DMFS) information, which allows us to specifically study the ability of EMAT clusters to predict the occurrence of metastasis. As can be seen in Figure 3A, the separation among Kaplan-Meier curves for the subtypes is statistically significant (p = 1.39E-2, log rank test). In addition univariable and multivariable Cox regression analysis (when considering clinical parameters) showed that these clusters provide a statistically significant prognostic value in this independent dataset as well which comprised only of LN patients who did not receive any adjuvant chemotherapy (Supplementary Table S5). It is interesting to note that the survival behavior of EMAT subtypes remains largely similar to their behavior in the LN METABRIC dataset, with EMAT4 having the worst survival, EMAT3 the second worst survival and EMAT1 and EMAT2 having the best survival probabilities. (We also tried a 5-nearest neighbor classifier for EMAT subtype status assignment which resulted in clusters with survival behaviors better matching those of Figure 1C; see Supplementary Figure S2.) Expression of the four biomarkers in GSE11121 samples assigned to each EMAT subtype generally follows a similar trend as in Figure 1C. These results show a high concordance between the characteristics of the EMAT subtypes in two independent studies.

**Figure 3:**
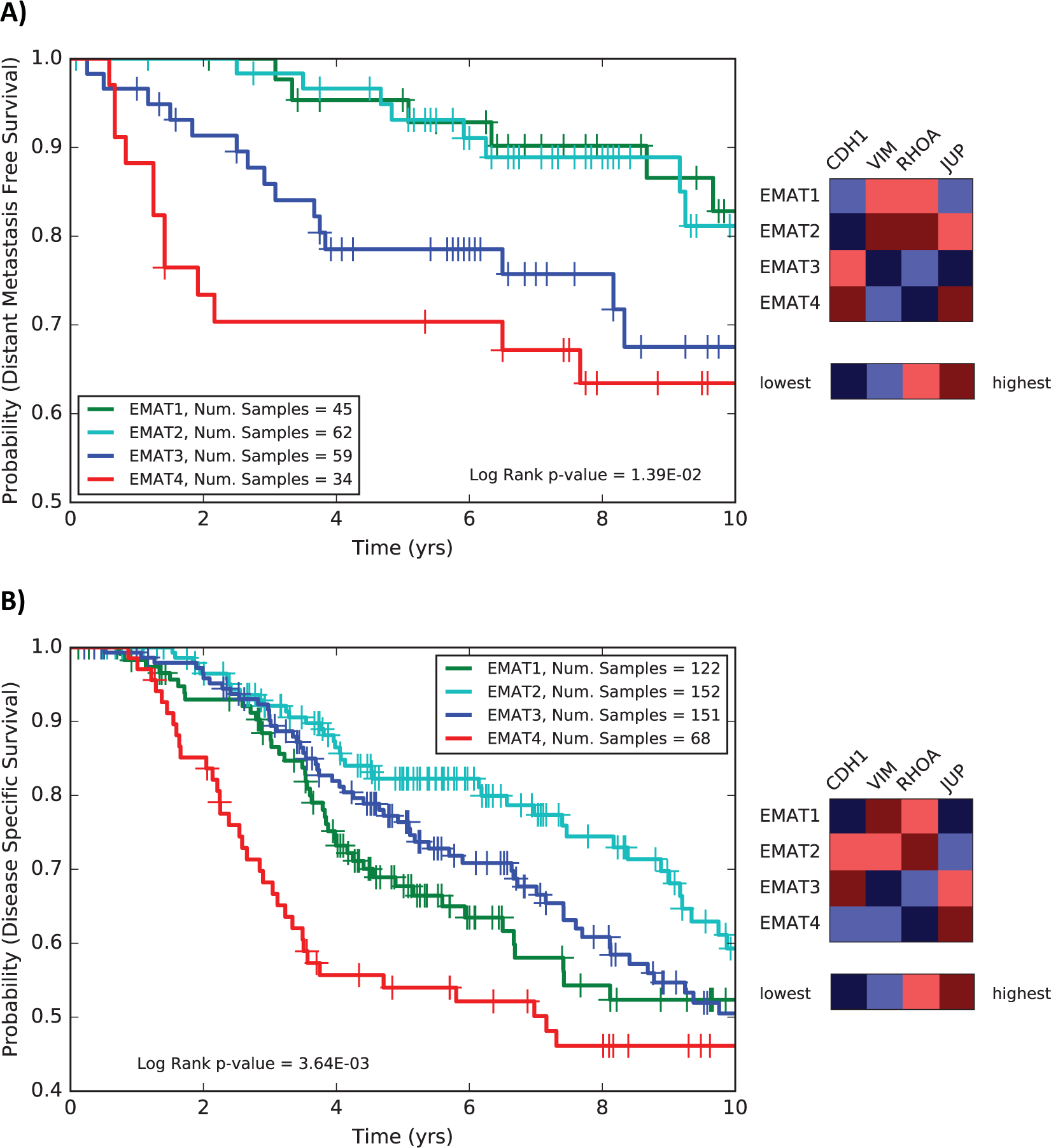
Cross-dataset analysis. A centroid-based classifier trained on LN METABRIC samples is used to assign EMAT subtype labels to each sample. (A) The Kaplan-Meier survival plots and biomarker status for EMAT subtypes of LN breast cancer samples from the GSE11121 dataset. Expression of the four biomarkers generally follows a similar trend as in Figure 1C; however, VIM is no longer highly expressed in EMAT4, but instead has the highest expression in EMAT2. In addition, expression of JUP is lowest in EMAT3, while it was highest in this cluster in Figure 1C (B) The Kaplan-Meier survival plots and biomarker status for EMAT clusters of LP breast cancer samples from the METABRIC dataset. The expression of biomarkers is largely consistent with LN samples. Two exceptions are the higher expression of VIM and CDH1 in EMAT2 and lower expression of VIM in EMAT4 in LP samples compared to LN samples in Figure 1C.

Up until now, by focusing on the LN samples, our analysis presumably reflected the metastatic mechanisms active prior to lymph node invasion. However, samples with similar gene expression patterns may have different clinical outcome conditioned on their lymph node status, as cells that have already acquired invasive characteristics may rely on different mechanisms to promote metastasis. To evaluate this, we obtained clinical and gene expression data corresponding to 493 lymph node-positive (LP) samples from the METABRIC study, and used our subtyping procedure to assign EMAT subtype designations to each sample. Kaplan-Meier analysis (Figure 3B) shows a significant distinction among survival probabilities of each EMAT subtype in LP patients (p = 3.64E-3). In addition, univariable and multivariable Cox regression analysis (when considering clinical parameters) showed that these subtypes provide a statistically significant prognostic value (Supplementary Table S3).

The main difference between the results obtained using the LP and LN samples is in the survival characteristics of EMAT1: while this subtype had the best survival probability in LN patients, it exhibits poor prognosis (second worst) in LP samples. However, the relative behaviors of EMAT2, EMAT3, and EMAT4 samples (Figure 3B) mimic those of their LN counterparts, in terms of their relative survival probabilities (Figure 1C) and biomarker characteristics (Figure 1C, inset). In summary, our analysis suggests that the clinical and molecular characteristics of the subtypes are preserved between LN and LP samples, with the exception of EMAT1: the EMAT1 phenotype when manifested prior to lymph node invasion, represents a less aggressive mode of cancer, but at later stages the same phenotype represents an aggressive mode, although biomarker expressions, enrichment in breast cancer subtypes, and resemblance to hESCs was similar in both LN and LP groups (Supplementary Figure S3). In the absence of nodal involvement, EMAT1 tumors appear to lack metastatic propensity, while the metastatic propensity of EMAT2, EMAT3 or EMAT4 tumors is manifested even in lymph node-negative patients. This suggests that the current classification is not adequate to capture the biologic heterogeneity of the EMAT1 phenotype. Future studies that can further sub-classify the EMAT1 phenotype may shed light as to why nodal involvement is not only predictive of an increased risk of metastatic dissemination in some patients, but may also likely to be a necessary mechanistic prerequisite for distant metastasis to occur.

### Identification of transcription factors associated with EMAT clusters

We evaluated the expression of 1,338 transcription factors (TFs) present in the METABRIC dataset. For this purpose, we used a t-test (corrected for multiple hypothesis testing) to identify TFs that are differentially expressed in one cluster compared to others, in LN samples (Supplementary Table S6). Next, for each cluster we identified two TFs that were most over-expressed and under-expressed (total of 8 TFs). These TFs include PPP1R13L and MNDA (EMAT1), ETV7 and TSHZ3 (EMAT2), SLUG and AFF3 (EMAT3), and FOXA1 and FOXC1 (EMAT4), which were under-expressed and over-expressed in each cluster, respectively, indicating a link with progression and invasion mechanisms (see Discussion). Hierarchical clustering of LN samples from the METABRIC dataset using the eight identified TFs had a very high concordance with clusters obtained using the full EMAT gene signature. In addition, Kaplan-Meier analysis using the clusters based on these TFs demonstrated a significant separation of survival curves (Figure 4). These results further support the key role of these TFs in potentially regulating the distinct expression profiles of EMAT subtypes and lend support to their clinical utility as potential biomarkers of the identified EMAT clusters.

**Figure 4:**
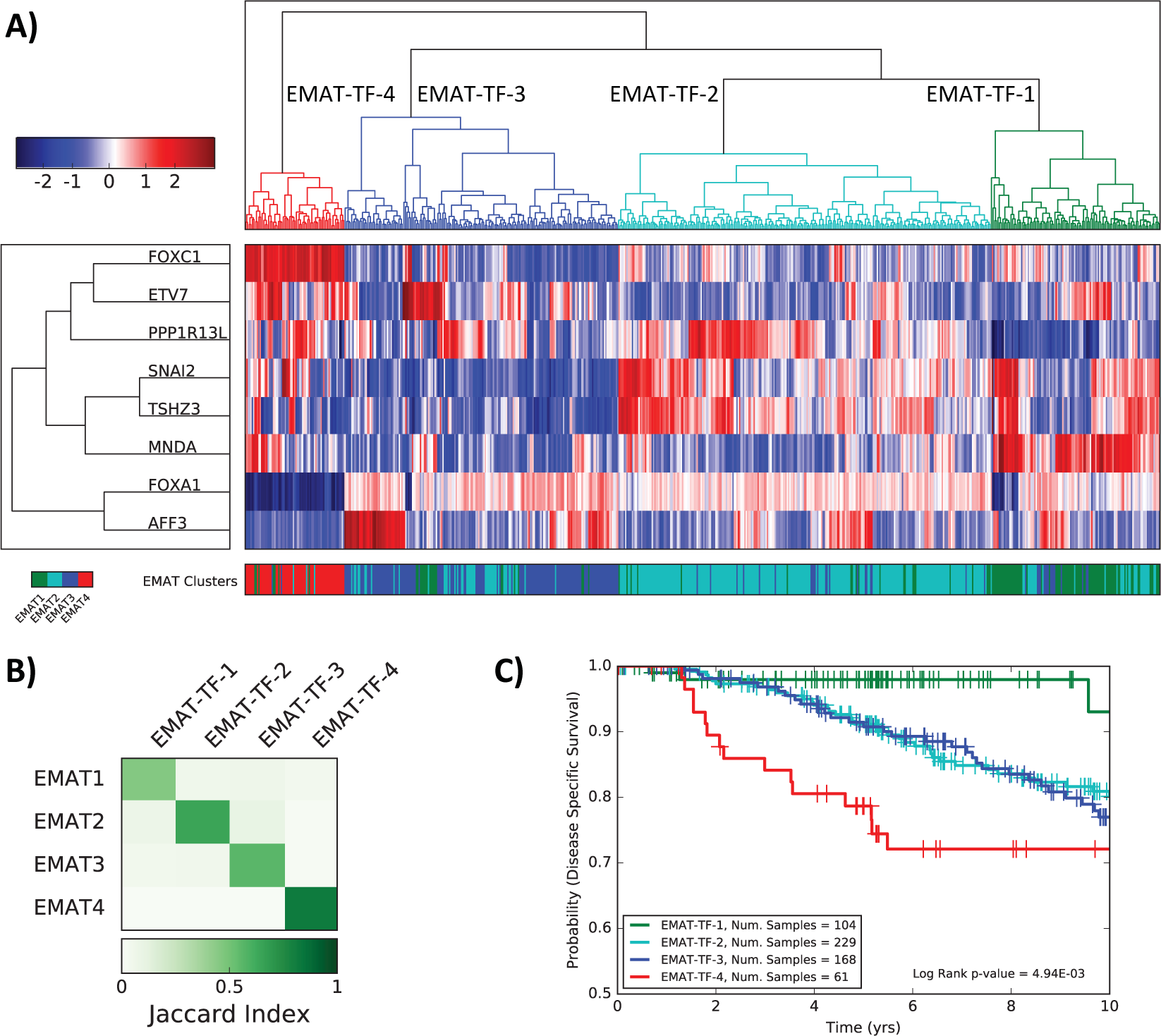
Analysis of clusters obtained using eight TFs most under-expressed or over-expressed in each EMAT cluster. (A) Hierarchical clustering based on the expression of the eight identified TFs is used to cluster samples into four groups. The color bar at the bottom shows the true EMAT cluster label of each sample. (B) Concordance of clusters obtained using eight TFs with EMAT clusters based on the Jaccard index. (C) Kaplan-Meier survival plots for clusters obtained using the TFs.

### EMAT subtypes reflect a progressive acquisition of malignancy and invasiveness

We next assessed the EMAT signature in the well-characterized isogenic MCF10 model of breast cancer progression which was produced by consecutive *in vivo* implantation of HRAS transformed MCF10A cells in immune-deficient mice followed by *in vitro* cell culture (Santner, Dawson et al., 2001). In a recent study that utilized whole genome, exome and RNA-Seq profiling of this isogenic MCF10A cell line series grown in 3D spheroid culture, investigators successfully identified genomic changes in these cell lines that were correctly representative of the stages of breast cancer progression, validating this system as a molecularly accurate model of breast cancer progression (Maguire, Peck et al., 2016).

RNA-Seq data corresponding to seven MCF10A-derived cell lines grown in three-dimensional (3D) spheroid culture systems (Maguire, Peck et al., 2016) were analyzed. For our analysis, we considered the cell lines in three groups - normal, benign and premalignant cells (MCF10A, MCF10AneoT, MCF10AT1), pre-invasive *in situ* carcinoma cells (MCF10DCIS.com), and invasive carcinoma cells (MCF10CA1a, MCF10CA1d, and MCF10CA1h) - and assessed their similarity to LN METABRIC EMAT clusters. For this purpose, we used the expression of the TFs identified above, except for MNDA and TSHZ3 which were not expressed in the cell lines. Figure 5 shows similarity between the centroids of each of the three cell line groups and those of the EMAT clusters, measured by pairwise Spearman’s rank correlation. Normal and pre-malignant cells were most similar to EMAT2, a cluster with relatively good survival compared to other clusters in both LN and LP samples. The non-invasive *in situ* cell line is most similar to EMAT3 cluster and the group containing invasive carcinoma cells is most similar to EMAT4, the cluster with the worst survival. Notably, the group containing invasive cell lines also has a high similarity to EMAT1 cluster. This supports the suggestion we made above, in the context of LP samples (Figure 3B), that the EMAT1 phenotype when manifested in later clinical stages of the invasion-metastasis cascade, i.e. post nodal involvement, denotes aggressive biology. Overall, these results support the hypothesis that the EMAT clusters identified *in vivo* represent a gradual progression of cancer cell transition from non-invasive to fully invasive states capable of metastasis. In addition, the observed association between the EMAT clusters with cell states of cancer progression in this model made intuitive sense with the progressively worsening prognosis of such EMAT clusters, as was observed in Figures 1C, 3A and 3B.

**Figure 5:**
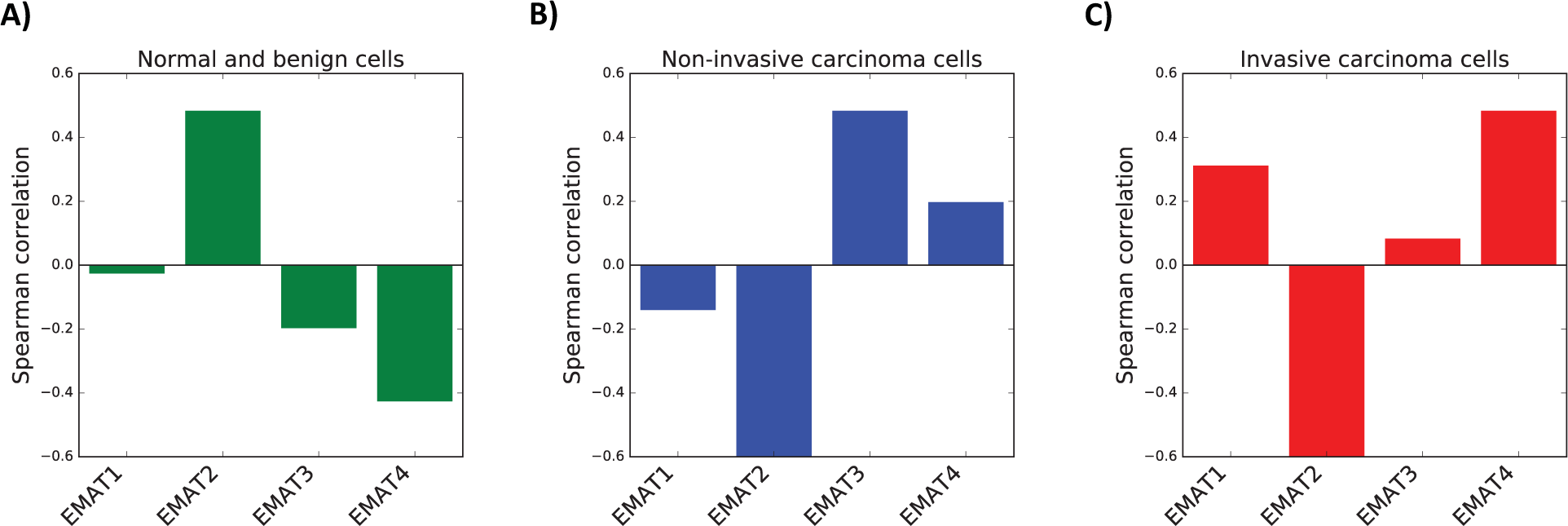
Analysis of MCF10 cell line series. (A) Spearman’s rank correlation of the centroid of normal and benign cells and EMAT clusters using expression of identified TFs. (B) Spearman’s rank correlation of the non-invasive carcinoma cells and EMAT clusters using expression of identified TFs. (C) Spearman’s rank correlation of the centroid of invasive carcinoma cells and EMAT clusters using expression of identified TFs.

## DISCUSSION

Prior attempts at utilizing EMT gene expression signatures of primary tumors to predict future metastasis have largely approached the problem as being able to define either one of two possible binary states, thereby treating metastasis as an “all-or-nothing” phenomenon (Tan et al., 2014, Taube et al., 2010). In these studies, it was assumed that EMT exists in either a turned “ON” or turned “OFF” state: this we thought might be too simplistic to discern the true survival impact of a complex tightly regulated biologic program like EMT, that most likely exists along an entire spectrum ranging from depressed states to elevated states. In our attempt to derive a gene expression signature based on metastasis biology that could classify this complex and dynamic process, we postulated that the heterogeneity that exists in the metastatic propensity of tumors might be better captured by considering the existence of multiple groups or clusters of patient tumors, each with characteristic molecular features and distinct prognostic profiles.

Of note, the clinical significance of EMT in driving metastasis has been questioned in recent publications that demonstrate the transient nature of EMT (Beerling, Seinstra et al., 2016) and dispute the necessity of EMT occurrence in mediating metastasis (Bill & Christofori, 2015, Diepenbruck & Christofori, 2016, Zheng, Carstens et al., 2015). While the findings in such studies have been challenged to be inconclusive (Aiello, Brabletz et al., 2017, Ye, Brabletz et al., 2017), there is another possible explanation for the reported findings without upending the potential significance of EMT in metastasis. One could speculate that when the cancer cell was proceeding along EMT, an increase in the entropy of an EMT path of cellular evolution could have resulted due to the cell being subjected to either microenvironmental pressures (e.g., hypoxia) or xenobiotic exposure (e.g., chemotherapy or EMT-targeted biologic therapy). A cancer cell of epithelial origin that had already undergone EMT when faced with such a situation could then undergo plastic transformation by a process such as MAT and adopt an alternate amoeboid motility program allowing it to proceed to metastasis via a lower entropy alternative route as recently reported (Lehmann et al., 2017). Indeed simultaneous targeting of both mesenchymal and amoeboid motility in an animal model of cancer progression has been demonstrated to effectively arrest metastatic spread (Jones et al., 2017). Only when both processes are considered together to contribute to multiple distinct molecular states of progressively increasing metastatic propensity does the prognostic value of such distinct phenotypes become demonstrable.

In this study, we utilized an EMAT gene expression signature and clustered breast cancer patients into four different groups with distinct prognosis, morphology, migratory behavior and molecular characteristics. Our results revealed the existence of subtypes of hybrid characteristics rather than discrete E-, M-, and A-like clusters, emphasizing the advantage of using the EMAT signature over using only E, M, or A biomarkers, to distinguish groups of patient tumors associated with distinct prognosis. Our findings suggest that during the invasion-metastasis cascade in cancer, both EMT and MAT contribute to an EMAT continuum of reverse evolution from a multicellular differentiated state towards a more primitive unicellular dedifferentiated state resembling embryonic stem cells (Chen, Lin et al., 2015). In addition, using *in vitro* cell line analysis, we showed that EMAT subtypes reflect a gradual and progressive acquisition of malignancy and invasiveness suggestive of the fact that EMAT subtypes with worse prognosis are more similar to hESC. The expression patterns of biomarkers of different modes of cell migration were generally consistent across different datasets. While in some subtypes one distinct morphology/motility mode was manifested, for other subtypes a hybrid of several morphologies was observed. Collectively, these results support the notion that several mechanisms are in effect simultaneously to promote metastasis, even among groups of individuals that share a similar transcriptomic profile. In addition, we showed that collective cell migration besides single cell migration is also a very important factor in determining clinical outcome, as individual tumors that exhibited elevated expression of the collective cell migration marker JUP consistently exhibited poor clinical outcome.

In order to investigate potential regulatory mechanisms underlying the discovered EMAT clusters, we identified transcriptional regulators characteristic of each EMAT cluster. In addition to confirming previously known factors, new factors involved in metastasis biology were also revealed. EMAT1 factor PPP1R13L, a known inhibitor of TP53, regulates apoptosis through NF-kappa-B and TP53 proteins and has been associated with lymph node metastasis in endometrial endometrioid adenocarcinoma. In addition, the over-expression of this gene has been shown to enhance tumorigenesis and invasion through both TP53-dependent and TP53-independent mechanisms (Laska, Lowe et al., 2009). MNDA, another predominant TF in the EMAT1 cluster, has been shown to be upregulated in tumorigenic breast cell lines (Roy, Calaf et al., 2001), and act as a master regulator in primary breast cancer (Baca-Lopez, Mayorga et al., 2012). ETV7, a factor implicated in the EMAT3 subtype, is known to be involved in cancer initiation, progression and metastasis (Sizemore, Pitarresi et al., 2017). TSHZ3, a TF which is mostly down regulated in the EMAT2 cluster, is a candidate tumor suppressor in breast and prostate cancers (Yamamoto, Cid et al., 2011). SLUG which is most upregulated in the EMAT3 cluster is a well-known EMT transcription factor (Taube et al., 2010), which represses E-cadherin (CDH1) expression in breast carcinoma. The role of this TF in metastasis is well documented in many cancers (Luanpitpong, Li et al., 2016, Zheng, Jiang et al., 2015). AFF3, another EMAT3 TF, has been shown to mediate the oncogenic effects of β-catenin in adrenocortical carcinoma (ACC) (Lefèvre, Omeiri et al., 2015). Finally, FOXC1 and FOXA1, the two TFs representing the cluster with worst prognosis (EMAT4), belong to the forkhead family of transcription factors. The roles of FOXC1 as an EMT driver and the opposing role of FOXA1 as an EMT suppressor are well established (Jensen, Ray et al., 2015, Ray, Jensen et al., 2015, Ray, Wang et al., 2010, Sizemore & Keri, 2012, Xu, Qin et al., 2017).

Our understanding of breast cancer progression and metastasis has largely been defined by three paradigms proposed over the past century. The first explanation conforms to the anatomic Tumor-Node-Metastasis model on which cancer staging is based. It proposed that breast cancer progression is an orderly process of contiguous spread, from primary tumor site, to regional lymph nodes via lymphatics, on to distant metastatic sites, and thus the anatomically precise removal of any and all demonstrable loco-regional disease, in the form of radical mastectomy, would favorably alter clinical outcome. This has come to be known as Halsted’s paradigm (Halsted, 1894). The second explanation, referred to as Fisher’s paradigm (Fisher, Bauer et al., 1985), was triggered by results of clinical trials proving the non-superiority of radical mastectomy, compared to lumpectomy and radiation. It proposed that breast cancer cells can spread and metastasize in a non-contiguous manner, wherein tumor biology trumps the anatomical extent of disease and small tumors are only a local manifestation of disease that has already disseminated. Hence their surgical removal would not favorably alter clinical outcome, as systemic disease already exists at the time of diagnosis. By the same token, nodal involvement is simply a clinical biomarker of distant disease already present and does not signify that adoption of the lymphatic route is mandatory in the path to distant metastasis. A third hypothesis accepts the existence of both of the above paradigms, and considers breast cancers to display a heterogeneous spectrum of metastatic proclivity, ranging from loco-regionally confined disease to systemically disseminated disease when first detected. This has come to be known as Hellman’s paradigm (Hellman, 1994, Hellman & Harris, 1987) and provides a rational explanation for why loco-regional treatment of some breast cancers with surgery and radiation is effective in favorably impacting clinical outcome, while seemingly ineffective at preventing distant metastasis in other cases.

In our study, the EMAT subtypes appear to support Hellman’s paradigm in that they support the existence of heterogeneity in the metastatic proclivity of breast cancers. What is important to note is that the metastatic propensity of breast cancer cannot be accurately captured or predicted by consideration of receptor status, PAM50 molecular subtype status or treatment variables alone, and that the additional consideration of metastasis biology is warranted. Our delineation of EMAT subtypes is an attempt to capture metastasis biology to improve prognostic prediction with regards to metastasis and further refinement of such approaches promises to increase the accuracy of prognostic prediction of breast cancer metastasis.

A key limitation of the present study is our inability to reconcile the prognostic impact of the EMAT1 subtype in LN and LP patients. As explained earlier, the current classification is not adequate to capture the biologic heterogeneity of the EMAT1 phenotype and future studies geared towards further sub-classifying the EMAT1 phenotype may shed light as to why nodal involvement is not only predictive of an increased risk of metastatic dissemination in some patients, but may also likely to be a necessary mechanistic prerequisite for distant metastasis to occur. Additional limitations of the present study include our inability to consider the potential impact of the host immune response, tumor stromal factors, and/or that of non-cancer cells of the tumor microenvironment on influencing metastatic proclivity. Future studies that are more comprehensive in collectively considering all potential factors that may biologically impact metastatic propensity of tumors hold promise of extending our understanding of metastasis biology and significantly improving prognostic prediction models.

Taken together, our analysis for the first time identifies EMAT subtypes of breast cancer progression and metastasis and provides a mathematically robust reasoning for the observed variance in metastatic propensity. While this advances our understanding of the heterogeneity and complexity of metastasis, delineation of the detailed molecular mechanisms underlying each EMAT subtype merits further investigation. Also, it still remains unclear whether the identified EMAT subtypes reflect the metastatic propensity of adenocarcinomas (originating from epithelial tissues) alone, or has an overlap with processes involved in driving metastatic propensity in cancers of mesenchymal origin as well.

## METHODS

### Derivation of MAT and EMAT gene lists

We obtained the list of 253 EMT-related genes from (Taube et al., 2010). This list had been derived in the original publication by analyzing gene expression data obtained from 5 distinct and separate EMT-inducing cell perturbation experiments to identify genes up- or down-regulated at least 2-fold in at least 3 experimental groups relative to control cells. Following identical methodology to minimize derivation bias, we derived a new MAT signature by analyzing gene expression data obtained from 4 distinct and separate MAT-inducing cell perturbation experiments (Taddei et al., 2014) to identify 138 genes up- or down-regulated at least 1.5-fold in at least 2 experimental groups relative to control cells. We then combined both the EMT and MAT gene signatures above to create the 388-member EMAT gene expression signature.

### Data Collection

We downloaded gene expression and clinical data corresponding to the METABRIC study from OASIS (http://oasis-genomics.org/) and supplemental material of (Curtis et al., 2012), respectively. In this dataset, 1055 samples (562 LN and 493 LP) had gene expression, survival information, and lymph node status. Gene expression and clinical information for cross-dataset analysis was downloaded from GEO (http://www.ncbi.nlm.nih.gov/geo/) under the accession number GSE11121, which contains 200 LN breast cancer samples. Gene expression profile of H1 hESC lines (Kim, Khalid et al., 2014) were downloaded from GEO under the accession number GSE54186. In all datasets, the probe intensities were log2 transformed and Z normalized prior to analysis.

### Hierarchical clustering and survival analysis

Hierarchical clustering of samples was performed using the python module SciPy, using Ward’s variance minimization algorithm. Survival analysis (Kaplan-Meier and Cox regression) was performed using lifelines python module (doi: 10.5281/zenodo.815943) and Survival R package (https://CRAN.R-project.org/package=survival). Variables included in the multivariate analysis are age, tumor size, and subtype status based on receptor profile, PAM50 centroid or EMAT centroid. All tests were two-sided, and P values of less than .05 were considered statistically significant.

### Cross validation framework to evaluate survival predictive ability of different predictors

In order to compare the predictive ability of different classes of features (Figure 2), we used a cross-validation framework, in which half of the samples (patients) were randomly selected as the training set and the other half were used as a test set. A Cox regression model was trained on the training set using one of the four options for predictor features described before, and the model was used to estimate the expected survival of each sample in the test set. Then, the estimated survival times were compared to the observed survival times using the concordance index (C-index). This process was repeated 200 times, each time using a distinct random partition of data. To determine the EMAT cluster assignment of the test samples, we first used hierarchical clustering to cluster samples in the training set into four clusters. Then we trained a centroid-based classifier on the training set and predicted the cluster labels for the test set.

### Cluster silhouette score calculations

The silhouette score (Rousseeuw, 1987) is a measure of the similarity of a sample to its own cluster compared to other clusters. A higher silhouette score averaged over all samples indicates a better separation of samples into clusters. We calculated this score for n = 3, 4, 5 clusters using Scikit-learn python module (http://scikit-learn.org). We used three metrics, the Euclidean, correlation and cosine distances. Two of these metrics showed that n = 4 clusters generate the highest average silhouette score, which we used for our analysis.

### The KNN and centroid-based classifiers

We implemented a KNN (only used to generate Supplementary Figure S2) and a centroid-based classifier for our analyses. In order to reduce the batch effects due to cross-dataset analysis and relax the normality assumption for gene expression values, we used the Spearman’s rank correlation as the measure of the similarity of samples in these classifiers. In the centroid-based classifier, for each cluster a centroid was calculated as a vector in which each entry corresponds to the mean expression of a gene across the samples of the cluster. The cluster whose centroid had the highest Spearman’s rank correlation with the expression profile of a test sample was selected as its label.

## Acknowledgements

We gratefully acknowledge and thank our friend and colleague Dr. Peter Friedl for his critical review of the manuscript and invaluable suggestions for improvement.

## Disclaimer

The study funders had no role in the design of the study, the collection, analysis, or interpretation of the data, the writing of the manuscript, nor the decision to submit the manuscript for publication.

## Authors’ Contributions

PSR and TR conceptualized and designed the study. AE, SS and PSR developed the methodology. AE, TR, TWJ, and PSR performed data acquisition. AE, TR, TWJ, MP, RN, SS and PSR analyzed and interpreted the data. AE, TR and PSR wrote the manuscript, which was edited by all the authors who approved the final version. SS and PSR provided study supervision. All authors are guarantors of the integrity of the data collection and interpretation.

